# Prototype-based AI triage for 3D pathology

**DOI:** 10.64898/2026.07.09.737559

**Authors:** Renao Yan, Gan Gao, Andrew H. Song, Huai-Ching Hsieh, Yujie Zhao, Cristina Almagro-Pérez, David Brenes, Sarah S.L. Chow, Jeanne Shen, Deepti M. Reddi, Lawrence D. True, Priti Lal, Anant Madabhushi, Faisal Mahmood, Jonathan T.C. Liu

## Abstract

Non-destructive 3D pathology enables high-resolution slide-free imaging of intact clinical specimens, providing comprehensive visualization of tissue structures beyond what conventional slide-based 2D histopathology can provide. However, the scale and complexity of volumetric datasets make exhaustive manual review impractical, motivating AI-assisted triage methods to select a small number of high-risk 2D slices for pathologist review. While prior triage models have shown promise, interpretability is poor and performance can be suboptimal, especially in the nascent field of 3D pathology in which labeled data is limited. We present SCOPE, a *S*egmentation-guided *C*r*O*ss-slice *P*rototyp*E* learning framework for comprehensive risk assessment of 2D levels within 3D pathology datasets. SCOPE combines (i) clustering-based pretraining on large-scale unlabeled volumetric data to initialize morphology-aware prototypes, (ii) segmentation-derived structural priors from publicly available models to guide prototype learning, and (iii) cross-slice (2.5D) prototype aggregation across neighboring slices to generate slice-level risk predictions. In prostate and esophageal data cohorts, SCOPE consistently outperforms attention-based and prototype-based multiple instance learning baselines for both binary and multiclass prediction tasks, enabling depth-resolved risk profiling for 3D triage based on morphological prototypes that are interpretable to pathologists.

## Introduction

Hematoxylin and eosin (H&E) slide-based histopathology is the clinical standard of care for most cancer diagnoses, underpinning clinical decision-making for millions of new cases per year ^1^. However, this centuries-old workflow examines only a small fraction of a clinical specimen (often *<*1% of the tissue volume) in the form of thin two-dimensional (2D) tissue cross-sections, which increases the risk of misinterpreting spatially heterogeneous, complex, and/or focal lesions. Moreover, the process is destructive and reduces the availability of valuable tissue for downstream molecular assays.

Recent advances in non-destructive three-dimensional (3D) pathology ^2–7^, such as with open-top light-sheet (OTLS) microscopy ^8–12^, enable high-resolution volumetric imaging of intact biopsies. By providing comprehensive interrogation of intact specimens without tissue destruction, including maintenance of molecular constituents ^13^, this paradigm has the potential to improve diagnostic performance and clinical workflows ^14–16^. However, the transition from 2D to 3D analysis introduces a challenge: the large size and structural complexity of 3D datasets make them impractical for pathologists to manually interpret in routine clinical practice ^17^ (**Figure 1a**).

**Figure 1:**
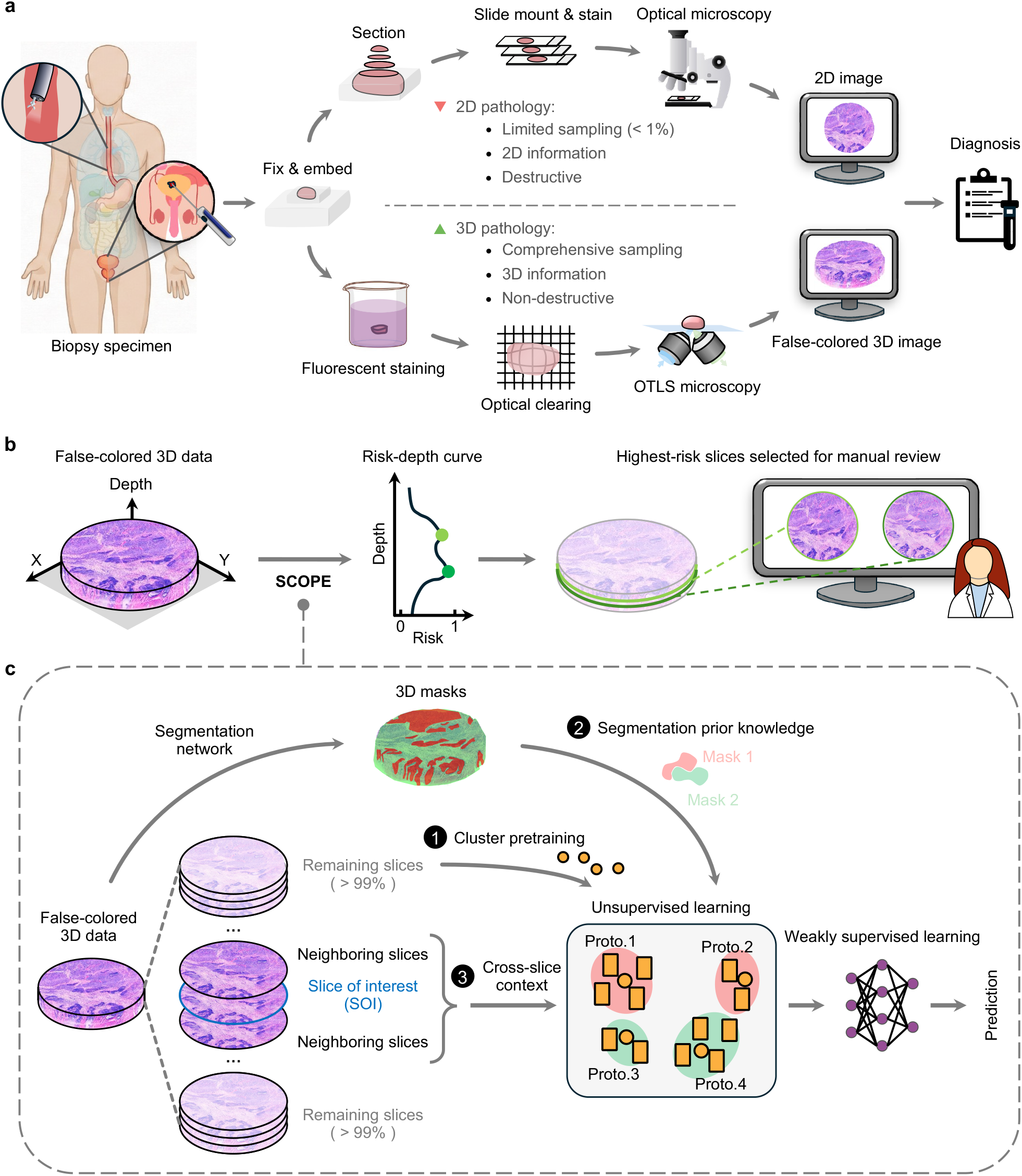
AI-triage of 3D pathology with SCOPE. (a) 2D pathology vs. 3D pathology. Standard slide-based histology requires destructive sectioning of tissue specimens to generate a few 2D sections from one side of the specimen, whereas 3D pathology enables nondestructive volumetric assessments of entire intact biopsies. **(b) AI-triaged 3D pathology**. Given a pathology volume, SCOPE assigns a risk score to each 2D slice and prioritizes a small set of the highest-risk slices for pathologist review. **(c) SCOPE components**. *(1) Cluster pretraining:* uses unlabeled cohort slices to initialize stable morphological prototypes. *(2) Segmentation priors:* uses segmentation masks to guide prototype learning toward biologically meaningful structures. *(3) Cross-slice context:* incorporates neighboring slices (2.5D) and a lightweight aggregator to improve slice-level risk predictions. visually recognizable tissue patterns ^25–28^. However, these approaches have been primarily developed for 2D pathology, in which out-of-plane tissue information within 3D pathology is not leveraged. In addition, the morphological prototypes are learned in an unsupervised manner, which can yield suboptimal or biologically ambiguous prototypes.

While fully automated end-to-end 3D classification pipelines provide one possible approach for analyzing volumetric pathology datasets ^18–20^, the lack of pathologist involvement in the interpretation process may hinder clinical adoption and regulatory approval. An alternative approach is to utilize AI to triage 3D pathology datasets and assist pathologists in rendering final diagnostic determinations ^21–23^. In this “AI-triaged 3D pathology” approach, a 3D dataset is treated as a stack of 2D slices, and the goal of the AI system is to assign a risk score to each slice within the stack based on predicted diagnostic relevance or disease severity. This enables a small number of high-risk slices to be prioritized for pathologists to review in a time-efficient manner. Ultimately, the clinical goal is to improve diagnostic assessment while maintaining or even reducing clinical workloads relative to standard-of-care 2D histopathology.

Weakly supervised learning is an attractive approach for developing AI models to assess the risk of 2D images because pathologists only need to provide diagnostic labels for entire 2D images rather than providing tedious pixel-level annotations. However, learning robust slice-level predictions is difficult within the relatively nascent field of 3D pathology, where only a limited number of 2D slices are annotated by pathologists ^24^. Furthermore, conventional weakly supervised learning approaches lead to AI models with limited interpretability. Prototype- or clustering-based learning addresses these challenges by compressing high-dimensional patch embeddings, many of which are highly redundant within tissues, into a small set of representative morphological prototypes ^25^. This reduces the effective solution space of the model and stabilizes weakly supervised learning. Beyond improved robustness, prototypes can promote interpretability by explaining predictions through

For AI-triaged 3D pathology, there are several opportunities to improve prototype learning. First, each volumetric specimen contains a large amount of unlabeled data (i.e., 2D image planes) containing substantial morphological diversity. This unlabeled 2D data can be leveraged to initialize the prototype-learning process, resulting in a stable set of morphological clusters. Second, high-performance 2D segmentation models now exist ^29–33^, as well as a small set of 3D segmentation approaches for specific tissue structures/compartments such as nuclei ^34^, glands ^18^, and tumor-enriched regions ^19^. While imperfect, these segmentation masks can serve as structural priors for prototype learning and can prevent biologically disparate morphologies from being clustered within the same prototypes. Finally, depth context from neighboring slices can be integrated for improved triage performance, consistent with routine clinical practice in which pathologists review multiple adjacent tissue levels to improve their diagnoses ^16^. This motivates a cross-slice aggregation strategy that integrates a small number of neighboring slices ^22, 23, 35^ while retaining the parameter efficiency needed for label-scarce settings.

In this study, we present SCOPE, a *S*egmentation-guided *C*r*O*ss-slice *P*rototyp*E* learning AI-triage framework designed for robust risk assessment when labeled 3D training data is limited (**Figure 1c**). In short, SCOPE is distinct from prior triage methods by integrating three components: (1) **unlabeled prototype pre-training**, which leverages the morphological diversity inherent in volumetric data to initialize stable prototype clusters; (2) **segmentation-derived structural priors** to guide these clusters to represent clinically and biologically relevant tissue structures; and (3) a **cross-slice (2.5D) prototype aggregator** that incorporates depth context from neighboring slices to improve the risk assessment of each slice of interest within the 3D dataset. Our results demonstrate that these design choices improve the robustness of AI triage for 3D pathology data. In addition, by enabling pathologists to inspect the morphological prototypes that drive slice-level risk scores, SCOPE provides a transparent and explainable approach for interpreting volumetric pathology datasets.

## Results

### Overview of the SCOPE framework

To bridge the gap between large-scale 3D pathology datasets and the practical constraints of pathologist-driven diagnostic determinations, we developed SCOPE: an AI-triaged 3D pathology framework that leverages morphological redundancy and 3D context to assign slice-level risk scores in a label-scarce setting.

SCOPE follows a weakly supervised learning paradigm (also referred to as multiple-instance learning, MIL) commonly used in computational pathology ^36–51^. Each 3D specimen is represented as a depth stack of 2D slices. Because exhaustive annotation across depth is impractical, only a small subset of slices are diagnostically labeled by pathologists. Each labeled slice is treated as a MIL “bag” composed of many image patches (“instances”) while the individual patches remain unlabeled. The model is therefore trained to summarize information across patches and predict the slice-level diagnostic category. In this study, we focus on multiclass risk assessment under limited supervision (small numbers of labeled image slices), a common situation in the nascent field of 3D pathology.

Given a slice of interest (SOI), SCOPE first divides tissue regions into patches and extracts embeddings using a fixed 2D feature encoder ^52, 53^. While attention-based aggregation is commonly used, learning stable attention weights over thousands of patch instances can be challenging, especially when only a small number of labeled slices are available. To mitigate this, SCOPE uses unsupervised prototype learning to compress a set of patch embeddings into a small, interpretable set of morphological prototype embeddings (**Extended Data Figure 1**). Specifically, patches with similar morphology are clustered into a limited number of learned prototypes (e.g., ~10), reducing thousands of patch embeddings into a concise set of prototype embeddings. This yields robust slice-level representations while improving the training stability in label-scarce settings. It also provides an interpretable link between model predictions and tissue morphology.

SCOPE adapts prototype-based MIL to 3D pathology through three design choices. First, it exploits unlabeled slices across the training cohort, beyond the limited set of pathologist-labeled slices, to initialize (pretrain) the prototype centroids. Second, it uses segmentation-derived structural priors to guide prototype learning, preventing biologically distinct structures from being merged into the same prototype. Third, it performs cross-slice aggregation to capture a limited range of neighboring depth context while keeping the aggregation lightweight and avoiding excessive parameterization. This helps to improve the model performance and minimize overfitting in label-scarce regimes.

SCOPE consists of two training phases: a cohort-level pretraining phase and a slice-level training phase (**Figure 2a**).

**Figure 2:**
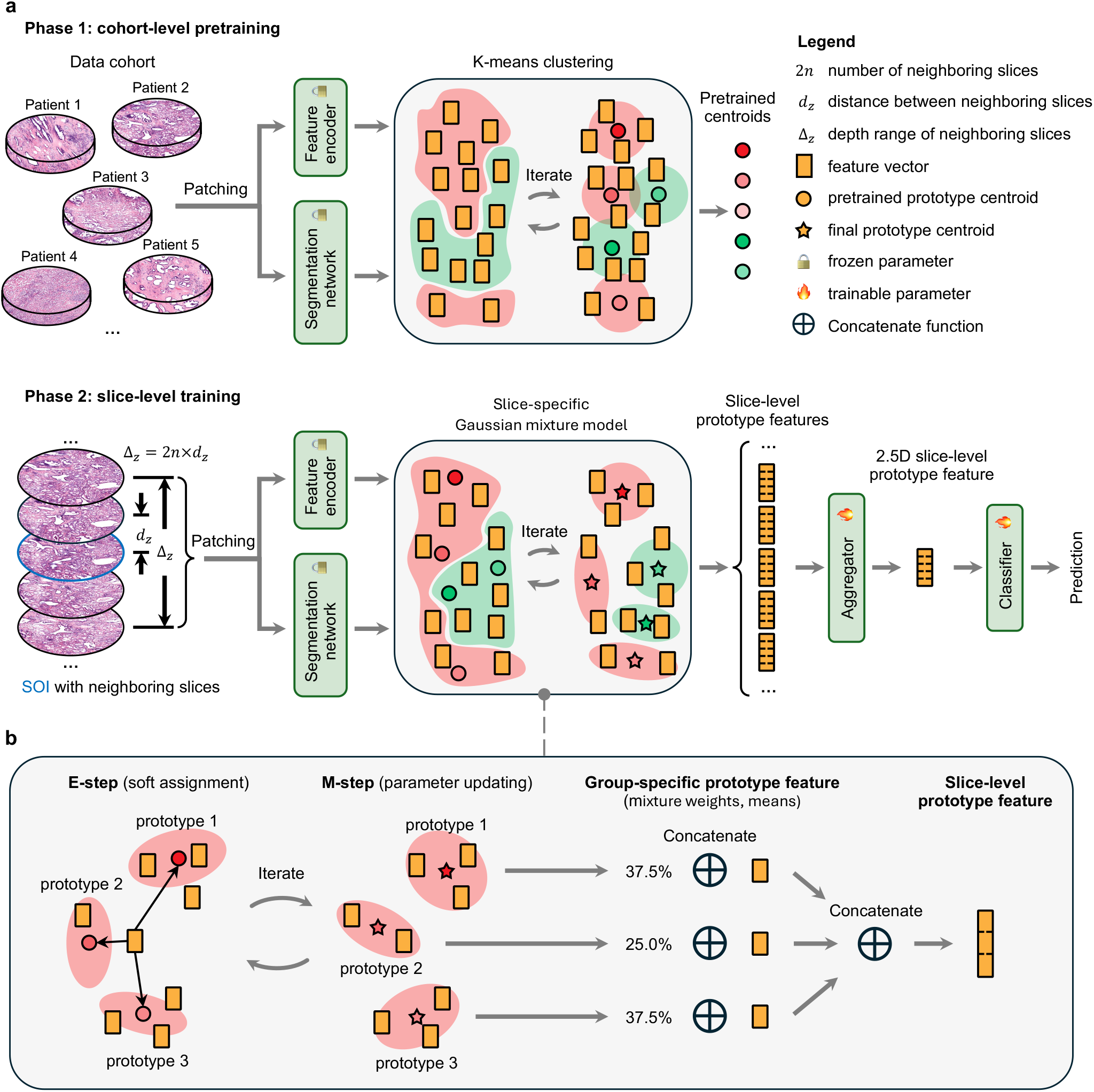
SCOPE framework overview. **(a)** Two-phase training with cross-slice (2.5D) inference. In Phase 1, patches from unlabeled cohort slices are encoded and clustered (K-means) within segmentation-defined tissue groups to initialize prototype centroids. In Phase 2, patches from a slice of interest (SOI) and neighboring slices are encoded and summarized within segmentation-defined tissue groups into prototype features using a Gaussian mixture model (GMM), with each mixture mean and weight representing a morphological concept and its prevalence within the slice, respectively. The prototype features are then fused by a lightweight cross-slice aggregator for slice-level risk prediction. **(b)** GMM-based prototype feature construction. Each mixture mean and weight is initialized to the corresponding prototype centroid and a uniform distribution, respectively. The expectation–maximization algorithm alternates between soft assignments (E-step) and parameter updates (M-step) to estimate slice-specific GMM parameters. These are concatenated across prototypes to form a fixed-length slice-level prototype feature. **Color code:** red and green indicate two segmentation-defined tissue types (e.g., glandular and non-glandular). Patches are assigned to a tissue type and prototypes are learned separately within each tissue type.

#### Phase 1: cohort-level pretraining

Patches from large unlabeled 3D pathology datasets are encoded using a fixed feature encoder, CONCH ^52^. Segmentation masks are then used to separate patches into tissue types. K-means clustering is performed on the patch features within each tissue type to initialize prototype centroids. In this work, we generate glandular and non-glandular masks for both prostate and esophageal tissues. For prostate, this is achieved by applying ITAS3D ^18^, a task-specific 3D gland segmentation method validated on OTLS-imaged prostate tissue, which provides consistent performance across depth. For esophagus, where task-specific 3D segmentation models are not yet available, we use a 2D segmentation foundation model, BiomedParse ^32^, to generate approximate gland masks. The resulting masks act as coarse structural priors rather than perfect annotations, and are used to steer prototype learning toward clinically and biologically relevant morphologies.

#### Phase 2: slice-level training

For each slice, CONCH-based patch embeddings within each segmentation-defined tissue group are clustered using a Gaussian mixture model (GMM). With a GMM, the set of patch embeddings is modeled as a mixture of Gaussian components. Each component is parameterized by the mean and variance of the patch embeddings within that cluster. In our application of GMM, the different components correspond to distinct morphological prototypes. The GMM also outputs the component weights, which indicate the prevalence of each prototype within a given tissue slice.

We apply GMM clustering to each slice in a 3D pathology dataset. For each slice, we initialize the component means with the pretrained cohort-level centroids from Phase 1 and initialize the component weights with a uniform distribution. We then fit the GMM using the expectation-maximization (EM) algorithm ^25, 54^ (**Figure 2b**). This yields slice-specific weights and means that summarize the components (morphological prototypes) within each slice. We concatenate these fitted weights and means (across all mixture components) to form the final slice-level prototype feature. We emphasize that the GMM clustering process is unsupervised and that only the downstream aggregator and classifier are trained during Phase 2 (**Figure 2a**).

To incorporate volumetric context during the downstream training, SCOPE extracts prototype features from the SOI and a limited number of neighboring slices. A cross-slice aggregator is then trained to fuse these features and generate an enhanced “2.5D representation” of the SOI. Because only a small number of neighboring slices are included (typically *<* 10), we use a lightweight attention module to aggregate depth context without introducing heavy parameterization. The aggregated representation is then passed through a linear classifier to produce the final slice-level risk score.

#### Inference

For predictions on unseen data, SCOPE performs a forward pass through the entire architecture. The goal is to assign a risk score to every depth slice within a 3D pathology dataset. For each SOI, we include a limited number of neighboring slices and compute slice-level prototype features using slice-specific

GMM clustering (initialized from the pretrained cohort-level centroids). The trained cross-slice aggregator then fuses the neighboring-slice prototype features with the SOI prototype features to create an enhanced “2.5D representation” of the SOI. Finally, a linear classifier is applied to obtain a slice-level risk score. This procedure is repeated over all slices in the volume to generate a depth-resolved risk profile that is used to prioritize slices for pathologist review.

### SCOPE achieves accurate risk assessment across diverse tissue cohorts

We benchmarked SCOPE against three representative MIL baselines: (i) ABMIL ^36^, a standard 2D attention-based MIL model that aggregates patch features within a single slice; (ii) PANTHER ^25^, a state-of-the-art 2D prototype-based MIL method that summarizes patch embeddings using a GMM-based set of morphological prototypes; and (iii) TRICARE ^22^, a 2.5D attention-based triage model that incorporates neighboring depth context. These baselines provide meaningful comparisons to isolate the contributions of prototyping (PANTHER vs. ABMIL) and cross-slice context (TRICARE vs. ABMIL), as well as to contextualize SCOPE as a segmentation-guided, cross-slice prototype-learning approach.

Evaluations were conducted using leave-one-patient-out cross-validation on three OTLS-imaged 3D pathology cohorts: two prostate cancer cohorts collected at the University of Washington (112 biopsy-core volumes from 54 patients; denoted Prostate) and the University of Pennsylvania (59 punch-biopsy volumes from 59 patients; denoted Prostate2), and one esophageal neoplasia cohort collected at the University of Washington (125 biopsy and EMR volumes from 29 patients; denoted Esophagus). Across all cohorts, we analyzed 142 patients and 296 biopsy volumes. Slice-level annotations are summarized as follows: Prostate, 121 annotated SOIs; Prostate2, 59 annotated SOIs; and Esophagus, 503 annotated SOIs (see Methods: Dataset description for details).

We evaluated the performance for multiclass risk assessments for two clinical applications. For prostate cancer, the four risk classes were based on Gleason Grade Group (GG): GG1, GG2, GG3, and GG4–5. For esophageal neoplasia screening, the four classes were non-dysplastic, low-grade dysplasia, high-grade dysplasia, and esophageal adenocarcinoma. Performance was primarily assessed using one-vs-rest area under the receiver operating characteristic curve (AUC), together with accuracy, precision, recall, and Cohen’s *κ*. In addition, because binary classification (low- vs. high-risk) is a common clinical threshold for guiding treatment decisions such as surveillance vs. curative therapy ^23^, we also report binary performance by grouping the lowest clinical category (GG1 for prostate; non-dysplastic for esophagus) versus all remaining categories. The statistical significance of SCOPE versus each baseline was assessed using a nonparametric bootstrap test with 5,000 resamples.

Across all three cohorts spanning prostate cancer risk assessment (Prostate and Prostate2) and esophageal neoplasia screening (Esophagus), SCOPE achieved the strongest overall performance compared with MIL baselines. This was true for both binary and multiclass classification (**Figure 3, Extended Figure 2**, and **3**). In the Prostate cohort (**Figure 3a**), SCOPE achieved AUCs of 0.907 (binary; 95% CI: 0.853–0.952) and 0.882 (multiclass; 95% CI: 0.829–0.926), improving over the best-performing baseline TRICARE (0.874/0.861; ΔAUC = 0.034/0.021) and showing significant gains over ABMIL and PANTHER. In the Esophagus cohort (**Figure 3g**), SCOPE achieved AUCs of 0.887 (binary; 95% CI: 0.852–0.919) and 0.889 (multiclass; 95% CI: 0.857–0.920), improving over TRICARE (0.872/0.872; ΔAUC = 0.015/0.017) and significantly outperforming ABMIL and PANTHER. Finally, in the Prostate2 cohort (**Extended Figure 3a**), SCOPE achieved AUCs of 0.873 (binary; 95% CI: 0.774–0.960) and 0.819 (multiclass; 95% CI: 0.707–0.924), significantly outperforming all baselines, with absolute improvements of ΔAUC = 0.137–0.147 (binary) and 0.120–0.132 (multiclass). Detailed performance comparisons are provided in **Extended Tables 1, 2, and 3**.

**Table 1:** Overall AUC performance with different encoders and MIL models on the Prostate dataset.

| **MIL Model** | **Encoder** | **Binary AUC** | **Multiclass AUC** |
| --- | --- | --- | --- |
| MeanMIL | CONCH | 0.794 | 0.785 |
| MaxMIL | CONCH | 0.761 | 0.763 |
| ABMIL | CONCH | 0.829 | 0.823 |
| CLAM | CONCH | 0.790 | 0.806 |
| DSMIL | CONCH | 0.784 | 0.795 |
| DTFD-MIL | CONCH | 0.790 | 0.789 |
| TransMIL | CONCH | 0.688 | 0.681 |
| RRT-MIL | CONCH | 0.813 | 0.820 |
| WIKG-MIL | CONCH | 0.787 | 0.773 |
| PANTHER | CONCH | 0.824 | 0.805 |
| TRICARE | CONCH | 0.873 | 0.861 |
| SCOPE | CONCH | 0.907 | 0.882 |
| MeanMIL | UNiv2 | 0.789 | 0.780 |
| MaxMIL | UNiv2 | 0.586 | 0.580 |
| ABMIL | UNiv2 | 0.856 | 0.805 |
| CLAM | UNiv2 | 0.845 | 0.807 |
| DSMIL | UNiv2 | 0.774 | 0.751 |
| DTFD-MIL | UNiv2 | 0.812 | 0.778 |
| TransMIL | UNiv2 | 0.772 | 0.731 |
| RRT-MIL | UNiv2 | 0.859 | 0.800 |
| WIKG-MIL | UNiv2 | 0.881 | 0.841 |
| PANTHER | UNiv2 | 0.884 | 0.841 |
| TRICARE | UNiv2 | 0.857 | 0.831 |
| SCOPE | UNiv2 | 0.892 | 0.846 |

**Table 2:** Overall AUC performance with different encoders and MIL models on the Prostate1 dataset.

| **MIL Model** | **Encoder** | **Binary AUC** | **Multiclass AUC** |
| --- | --- | --- | --- |
| MeanMIL | CONCH | 0.763 | 0.721 |
| MaxMIL | CONCH | 0.738 | 0.651 |
| ABMIL | CONCH | 0.729 | 0.687 |
| CLAM | CONCH | 0.769 | 0.709 |
| DSMIL | CONCH | 0.738 | 0.700 |
| DTFD-MIL | CONCH | 0.752 | 0.723 |
| TransMIL | CONCH | 0.685 | 0.646 |
| RRT-MIL | CONCH | 0.582 | 0.557 |
| WIKG-MIL | CONCH | 0.731 | 0.672 |
| PANTHER | CONCH | 0.727 | 0.699 |
| TRICARE | CONCH | 0.736 | 0.696 |
| SCOPE | CONCH | 0.873 | 0.819 |
| MeanMIL | UNiv2 | 0.751 | 0.695 |
| MaxMIL | UNiv2 | 0.720 | 0.602 |
| ABMIL | UNiv2 | 0.789 | 0.742 |
| CLAM | UNiv2 | 0.796 | 0.725 |
| DSMIL | UNiv2 | 0.750 | 0.703 |
| DTFD-MIL | UNiv2 | 0.832 | 0.776 |
| TransMIL | UNiv2 | 0.661 | 0.592 |
| RRT-MIL | UNiv2 | 0.755 | 0.686 |
| WIKG-MIL | UNiv2 | 0.773 | 0.745 |
| PANTHER | UNiv2 | 0.743 | 0.665 |
| TRICARE | UNiv2 | 0.764 | 0.705 |
| SCOPE | UNiv2 | 0.812 | 0.707 |

**Table 3:** Overall AUC performance with different encoders and MIL models on the Esophagus dataset.

| **MIL Model** | **Encoder** | **Binary AUC** | **Multiclass AUC** |
| --- | --- | --- | --- |
| MeanMIL | CONCH | 0.783 | 0.793 |
| MaxMIL | CONCH | 0.781 | 0.782 |
| ABMIL | CONCH | 0.858 | 0.853 |
| CLAM | CONCH | 0.832 | 0.833 |
| DSMIL | CONCH | 0.863 | 0.860 |
| DTFD-MIL | CONCH | 0.701 | 0.706 |
| TransMIL | CONCH | 0.743 | 0.734 |
| RRT-MIL | CONCH | 0.861 | 0.860 |
| WIKG-MIL | CONCH | 0.774 | 0.778 |
| PANTHER | CONCH | 0.851 | 0.854 |
| TRICARE | CONCH | 0.872 | 0.872 |
| SCOPE | CONCH | 0.887 | 0.889 |
| MeanMIL | UNIV2 | 0.792 | 0.798 |
| MaxMIL | UNIV2 | 0.721 | 0.713 |
| ABMIL | UNIV2 | 0.686 | 0.686 |
| CLAM | UNIV2 | 0.813 | 0.812 |
| DSMIL | UNIV2 | 0.778 | 0.779 |
| DTFD-MIL | UNIV2 | 0.762 | 0.766 |
| TransMIL | UNIV2 | 0.733 | 0.737 |
| RRT-MIL | UNIV2 | 0.808 | 0.805 |
| WIKG-MIL | UNIV2 | 0.735 | 0.742 |
| PANTHER | UNIV2 | 0.791 | 0.790 |
| TRICARE | UNIV2 | 0.811 | 0.807 |
| SCOPE | UNIV2 | 0.817 | 0.819 |

**Figure 3:**
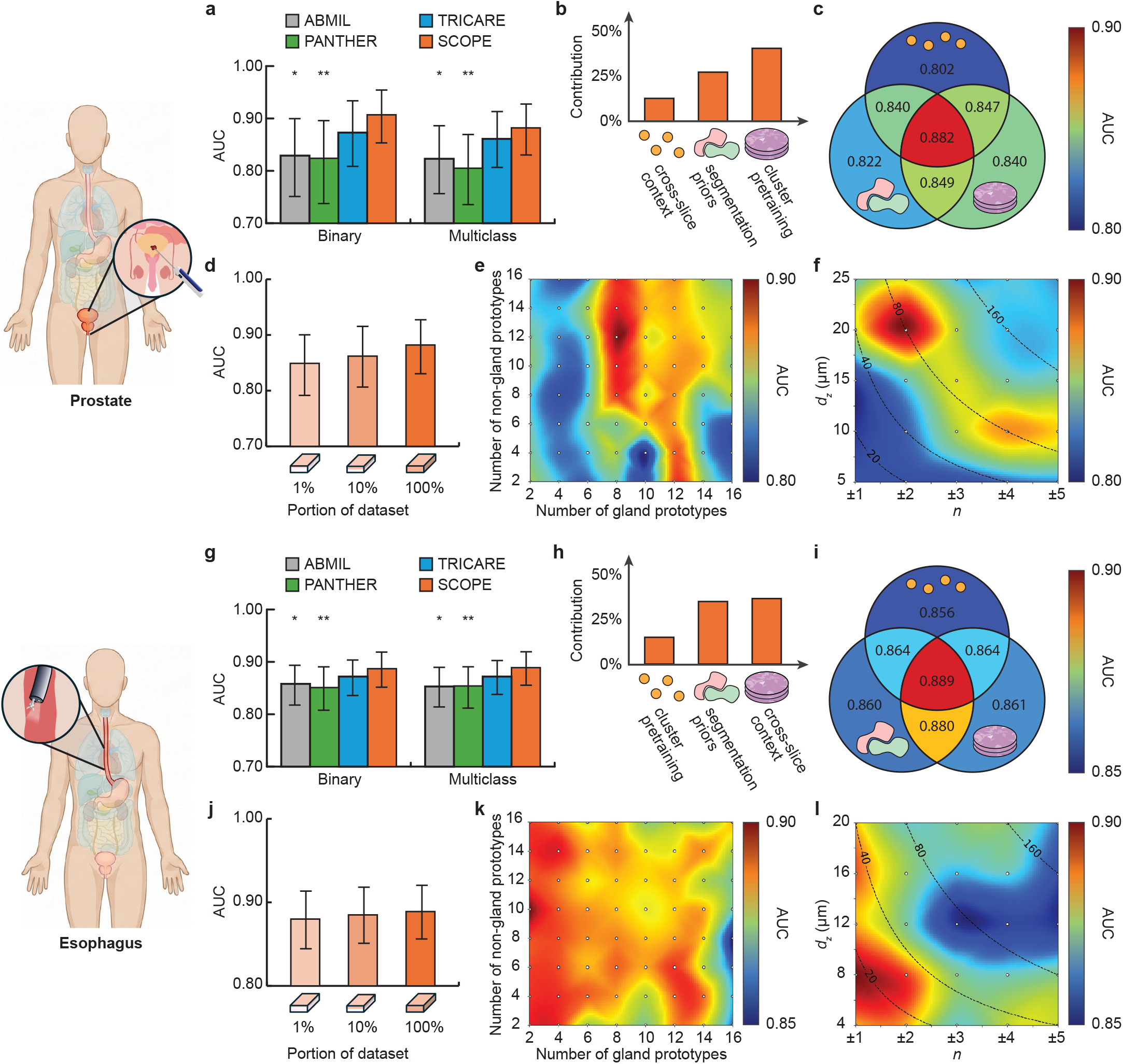
Risk assessment performance and ablation studies across cohorts. Results are shown for Prostate **(a–f)** and Esophagus **(g–l). (a, g)** Binary and multiclass AUC for SCOPE and baselines (ABMIL, PANTHER, TRICARE). **(b, h)** Shapley value (SHAP)-based contribution of the three design choices (prototype pretraining, segmentation priors, cross-slice context). **(c, i)** Performance of all 2^3^ component combinations (Venn diagrams; center = full SCOPE). **(d, j)** Effect of unlabeled pretraining scale on downstream AUC. **(e, k)** Sensitivity to the number of gland and non-gland prototypes. **(f, l)** Sensitivity to cross-slice context parameters (number of neighboring slices on each side *n* and spacing *d*_*z*_, with total depth context Δ_*z*_ = 2*n × d*_*z*_). Bars show mean AUC across leave-one-patient-out folds; error bars indicate 95% confidence intervals. Statistical significance is shown for SCOPE versus each baseline (^*^*p* ≤ 0.05, ^**^*p* ≤ 0.01; nonparametric bootstrap test with 5,000 resamples).

To understand the contribution of each design component, we performed two complementary analyses. First, we computed Shapley values (SHAP) ^55^ for the three main design choices (prototype pretraining, segmentation priors, and cross-slice context) to estimate the marginal contribution of each design component to the AUC (**Figure 3b, h**; **Extended Figure 3b**). Second, we performed a controlled component-combination study in which the components were individually enabled/disabled and all 2^3^ model variants were trained (**Figure 3c, i**; **Extended Figure 3c**).

Across cohorts, SHAP analysis indicates that both segmentation priors and cross-slice context contribute strongly, with the relative contribution of each component varying by dataset: cross-slice context contributes more for the primary Prostate and Esophagus cohorts (**Figure 3b, h**), whereas segmentation priors have a slightly greater impact on the Prostate2 cohort (**Extended Figure 3b**). Prototype pretraining provides a smaller but consistent benefit. The best performance is achieved only when all three components are enabled, indicating that they provide complementary benefits (**Figure 3c, i**).

We further analyzed several hyperparameter configurations within the three components that affected downstream performance:

#### Pretraining scale

Increasing the amount of unlabeled 3D data used for centroid initialization consistently improves performance across cohorts, suggesting that larger pretraining sets yield more representative prototype centroids (**Figure 3d, j**; **Extended Figure 3d**). To construct reduced pretraining sets (e.g., 1% and 10%), we uniformly subsampled the unlabeled slices along the axial (*z*) dimension across the training cohort.

#### Clustering architecture

A grid search over the number of clusters indicates that gland-related clusters are most influential for diagnosis, with the optimal number being 8 gland clusters for prostate and 2 gland clusters for esophagus (**Figure 3e, k**; **Extended Figure 3e**). In contrast, the optimal number of non-gland clusters varies across cohorts. This reflects clinical practice, in which glandular structures are currently relied upon for prognostic Gleason grading whereas cytologic features are more important for diagnosing/grading esophageal lesions.

#### Spatial configuration

For prostate, optimal performance is obtained with a total cross-slice depth context of ~80 *µ*m, with two neighboring slices on each side of the SOI in one cohort (**Figure 3f**) and three on each side in the other (**Extended Figure 3f**). For esophagus, the optimal performance is achieved with a total depth context of ~20 *µ*m with one slice above and below the SOI (**Figure 3l**). These findings align with clinical practice: prognostic Gleason grading relies on large glandular morphologies in which a larger depth context is helpful, whereas the diagnosis and grading of esophageal lesions relies more on finer cytologic features in which a smaller depth context is preferable.

### Segmentation guidance generates biologically meaningful prototypes

To evaluate whether segmentation-guided priors enhance the biological relevance of morphological prototypes, we analyzed the initial prototypes learned during cohort-level pretraining in the Prostate cohort. For prostate gland segmentation, we used ITAS3D ^18^, a task-specific 3D gland segmentation model developed and validated for OTLS-imaged prostate tissue. In contrast, for esophagus, there is currently no task-specific 3D segmentation model; therefore, we used the general-purpose foundation model BiomedParse ^32^ to generate approximate gland masks. Since more-reliable 3D gland-segmentation priors are available for prostate cancer (rather than esophagus), we focused the following analyses on the Prostate cohort.

We compared conventional (non-segmentation-based) clustering versus segmentation-guided clustering. For conventional clustering, patch embeddings from the entire Prostate cohort were pooled and clustered directly (K-means, *K* = 6, chosen via validation). By contrast, SCOPE first partitioned embeddings into glandular and non-glandular sets using segmentation-derived labels and then clustered the patches within each set.

To visualize the learned prototypes, we projected patch representations and prototype centroids into two dimensions using principal component analysis (PCA) (**Figure 4a**). Qualitatively, the conventional clustering approach shows regions in which glandular and non-glandular embeddings are mixed, suggesting that purely appearance-based clustering can mix biologically distinct tissue types. To quantify this effect, we computed for each conventional cluster the fraction of patches assigned as glandular versus non-glandular by the segmentation model. The hierarchical pie chart in **Figure 4b** shows that multiple conventional clusters (e.g., *c*2–*c*5) contain substantial contributions from both tissue types, indicating limited semantic specificity and biological relevance in the absence of segmentation guidance.

**Figure 4:**
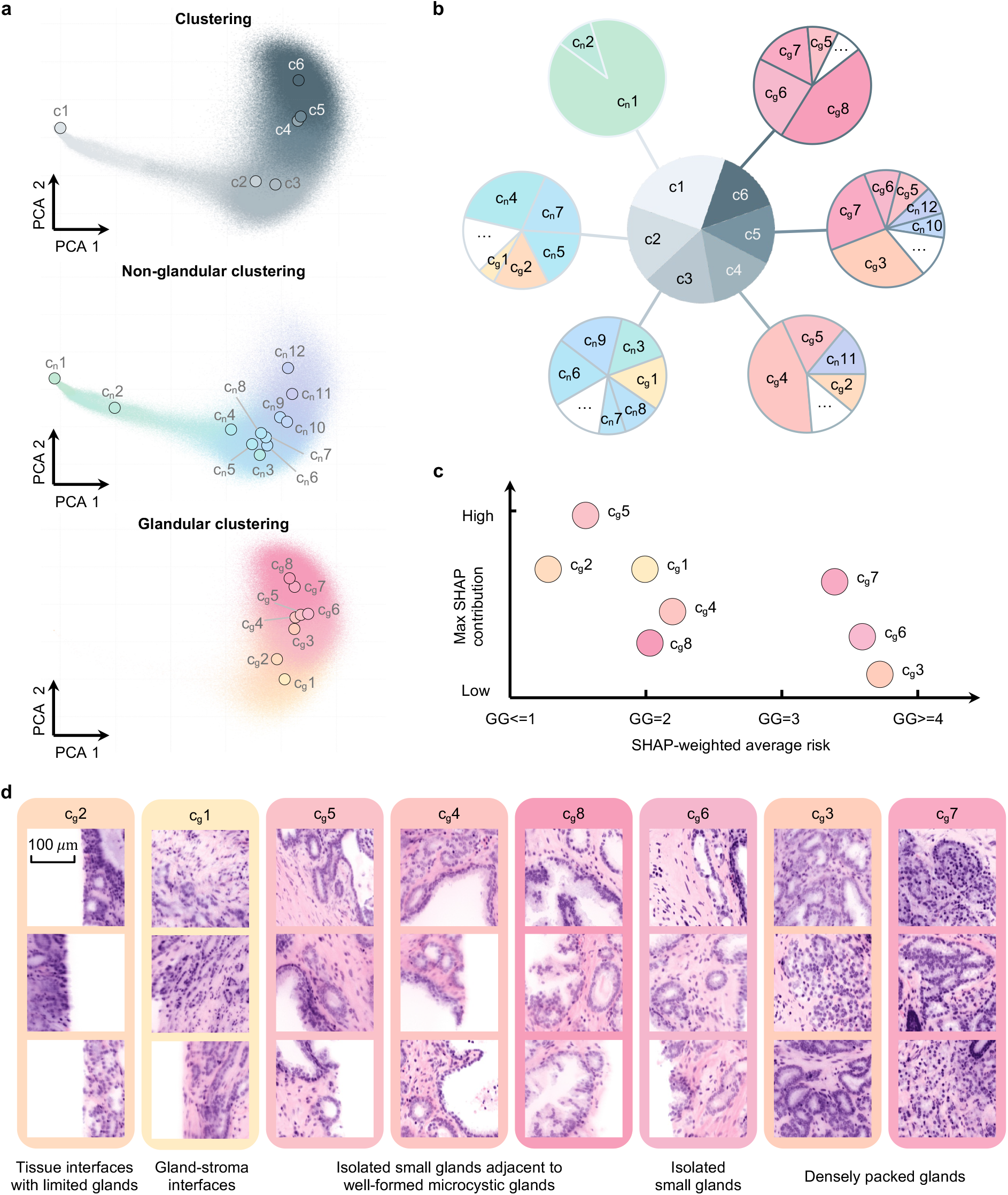
Interpretability of segmentation-guided clustering in the Prostate cohort. **(a)** PCA projection of patch embeddings (points) and K-means centroids (circles) for conventional clustering (top), non-glandular clustering (middle), and glandular clustering (bottom). **(b)** Center pie shows conventional prototypes; outer pies show the corresponding segmentation-guided prototypes, revealing that conventional prototypes often contain a mixture of glandular and non-glandular morphologies with reduced semantic specificity and biological relevance. **(c)** SHAP-based contribution analysis for glandular prototypes. **(d)** Representative patches nearest to each glandular centroid, reviewed by pathologists and consolidated into coarse morphologic categories. Gray, cool (blue), and warm (pink) colors indicate conventional, non-glandular and glandular prototypes, respectively. Scale bar represents 100 *µ*m.

We next investigated the morphological and clinical relevance of the glandular prototypes learned by SCOPE. As shown in **Extended Figure 4**, we computed SHAP values to quantify prototype-level contributions to the risk prediction. Using these attributions, we summarized each glandular prototype’s association with predicted risk by reporting a SHAP-weighted average risk score, together with the maximum absolute SHAP contribution (**Figure 4c**).

Prototype *c*_*g*2_ is primarily associated with the low-grade disease and exhibits limited influence on the risk prediction. Prototypes *c*_*g*1_, *c*_*g*4_, *c*_*g*5_, and *c*_*g*8_ are enriched in grade group 2 (GG2) predictions and show relatively large SHAP contributions, indicating that they frequently contribute to model decisions. Prototypes *c*_*g*6_ and *c*_*g*7_ are associated with higher-grade diseases (GG3–GG≥4) and retain relatively large SHAP contributions, suggesting that these morphologies serve as strong indicators of the aggressive disease. Prototype *c*_*g*3_ exhibits the highest SHAP-weighted average risk but the lowest maximum SHAP contribution. This may reflect a morphology that is less prevalent across the cohort or contributes to model predictions in conjunction with other morphologies.

We also computed SHAP scores for non-glandular prototypes (Extended Figure 4b). As these prototypes do not show strong, consistent positive or negative associations with risk, their contributions are shown in gray. This is consistent with clinical practice, where Gleason grading primarily relies on glandular architecture. Therefore, we focus subsequent qualitative visualizations on the glandular prototypes.

As a qualitative validation of the learned glandular prototypes, we compiled three representative patches per prototype (nearest to each centroid) and had them reviewed by pathologists. Based on shared histomorphologic patterns, the eight glandular prototypes were organized into five coarse morphologic categories (**Figure 4d** and **Supplementary Video 1**). Prototype *c*_*g*2_ consists primarily of tissue interfaces with limited glandular content. Gland–stroma interfaces are mostly captured by *c*_*g*1_, characterized by stroma-dominant areas with sparse glandular structures. A third grouping comprising *c*_*g*5_, *c*_*g*4_, and *c*_*g*8_ is characterized by isolated small glands adjacent to well-formed larger glands, whereas *c*_*g*6_ captures small glands in a more isolated configuration. Finally, *c*_*g*3_ and *c*_*g*7_ consist mostly of densely packed glands, encompassing high-risk morphologies such as poorly formed glands, fused glands, and cribriform patterns.

### SCOPE generates interpretable depth-resolved risk profiles

A key advantage of SCOPE is that it moves beyond providing a single prediction per specimen and instead offers a spatially resolved view of disease risk across depth. This enables pathologists to identify where diagnostically important patterns appear within an intact 3D biopsy, including regions that may be missed by sparse 2D sampling. To support clinical decision-making, SCOPE can be used to generate: (i) the predicted diagnostic category at each depth level and (ii) a risk score at each depth level. To generate these results, SCOPE is first applied to every depth slice within a 3D biopsy volume to produce a diagnostic prediction (binary or multiclass). A depth-resolved risk profile is generated by plotting the confidence score of the highest-risk category detected within each specimen as a function of depth.

In a representative prostate biopsy (**Figure 5a** and **Supplementary Video 2**), SCOPE predicts GG1 at 358 *µ*m, consistent with the presence of numerous well-formed glands (ROIs 1–4). In contrast, at 90 *µ*m the model predicts GG2, driven by regions containing abundant fused glands (ROI 5), despite the presence of some well-formed glands (ROI 6). For the esophagus example (**Figure 5b**), SCOPE predicts esophageal adenocarcinoma at 619 *µ*m, supported by stromal invasion (ROI 7), whereas at 309 *µ*m it predicts low-grade dysplasia, characterized by nuclear hyperchromasia and pseudostratification without loss of polarity (ROI 8). These examples illustrate that depth-resolved multiclass predictions can prioritize slices not only by predicted probabilities, but also by clinically ordered severity.

**Figure 5:**
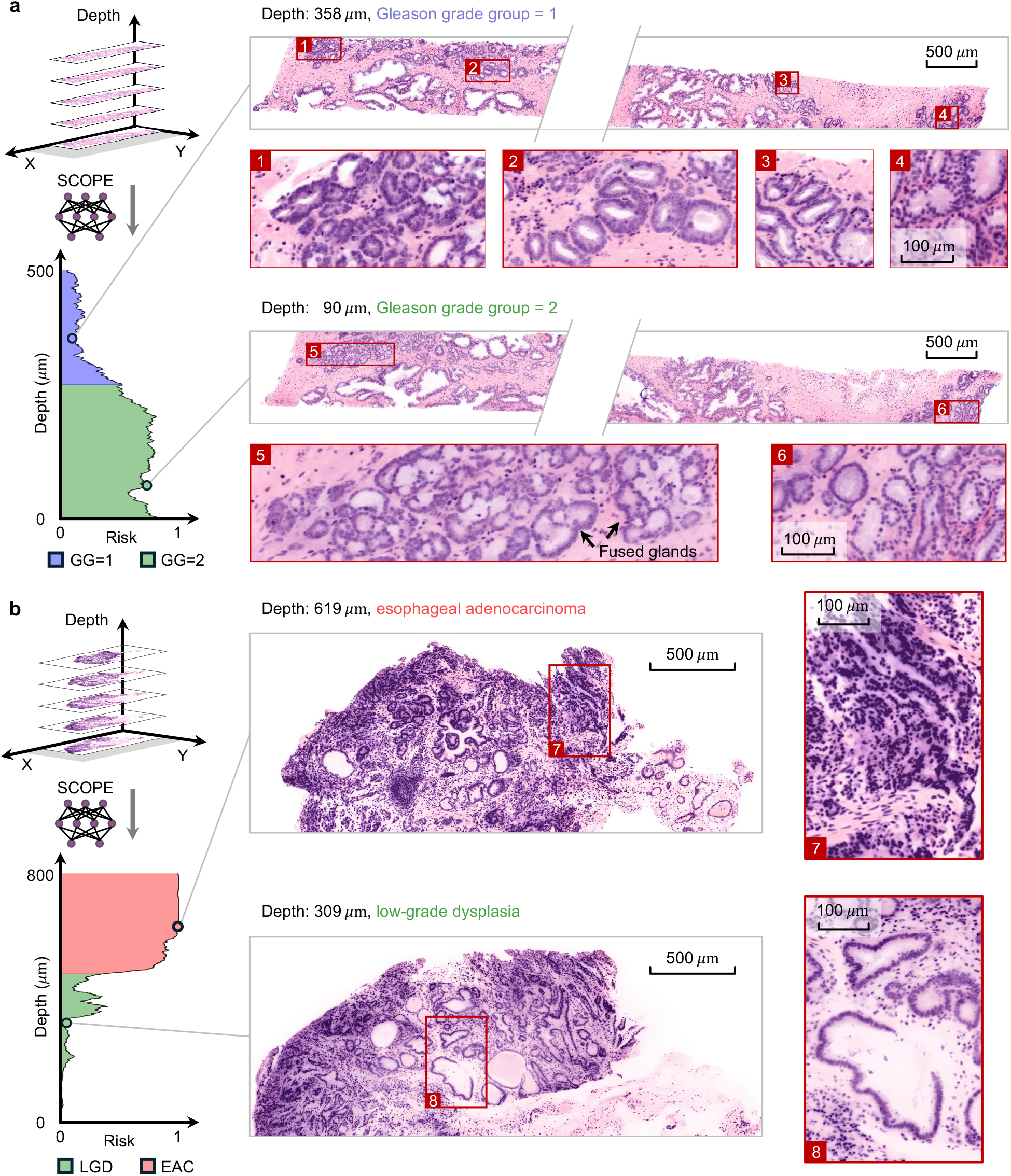
Interpretable 3D triage via risk–depth curves in representative (a) prostate and (b) esophageal biopsies. Left panels show risk–depth curve generated by SCOPE across all depth slices; colors indicate the predicted class at each depth, and the curve height indicates the risk score (probability of the specimen-level target risk class). Right panels show example whole-slice views at selected depths, with red boxes marking ROIs shown at higher magnification. Representative slices of different categories illustrate depth-dependent heterogeneity. Scale bars represent 500 *µ*m for whole-slice views, and 100 *µ*m for ROI zoom-ins.

We further highlight a representative 3-mm punch biopsy from the Prostate2 cohort in **Extended Figure 5**. Using SCOPE’s risk–depth curve for AI triage, the top-ranked high-risk slices (e.g., 385 *µ*m and 159 *µ*m) are selected for pathologist review. Both slices contain poorly formed glands, but they capture complementary high-risk morphologies: glomeruloid glands are identified at 385 *µ*m, whereas cribriform pattern is captured at 159 *µ*m.

Collectively, these results demonstrate that SCOPE can decompose complex 3D pathology volumes into interpretable depth-resolved risk profiles. By prioritizing diagnostically relevant slices across depth, SCOPE enables time-efficient review of heterogeneous specimens that may be undersampled and misdiagnosed with conventional 2D histopathology.

## Discussion

Non-destructive 3D pathology enables comprehensive visualization of intact clinical specimens, but it also introduces a major practical challenge: the scale of volumetric data makes exhaustive manual review infeasible within routine clinical workflows. While end-to-end 3D deep learning models can provide volume-level predictions, clinical adoption and regulatory approvals may be hindered by their limited interpretability and/or lack of pathologist oversight. AI-assisted triage potentially offers a more clinically deployable alternative by prioritizing a small set of 2D image slices for targeted assessment, allowing pathologists to remain responsible for final diagnostic determinations. However, due to a lack of large cohorts of human-annotated images from 3D pathology datasets, it is challenging to train AI-triage models, which are often highly parameterized and at risk of overfitting.

Motivated by the need for a high-performance AI-triaged 3D pathology pipeline that can be trained with limited labeled data, we developed SCOPE. SCOPE produces depth-resolved risk profiles based on binary or multiclass predictions, enabling pathologists to prioritize the most representative levels within a 3D whole-specimen dataset. SCOPE is grounded in the fundamental concept of morphological prototyping, which reduces model complexity (parameter space) and thereby mitigates overfitting. These prototypes also provide an interpretable link between model prediction and the underlying tissue morphologies/prototypes that contribute to those predictions. Specifically, SCOPE integrates three complementary concepts: (1) clustering-based pretraining on large-scale unlabeled volumetric data to improve performance through better initialization of morphology-aware prototypes; (2) segmentation-derived structural priors to facilitate the generation of prototypes that are more aligned with biological understanding; and (3) cross-slice integration to leverage morphological context across a limited number and range of neighboring depth levels. For prostate cancer grading and esophageal dysplasia/cancer screening, SCOPE consistently outperforms attention-based and prototype-based MIL baselines for both binary and multiclass diagnostic tasks. Importantly, ablation studies show that peak performance requires all three of the innovative components of SCOPE listed above.

As mentioned, a key practical advantage of SCOPE is increased interpretability and biological relevance. Segmentation-guided clustering reduces mixing between glandular and non-glandular patterns and produces prototypes that are aligned with recognizable histomorphologies (e.g., densely packed/cribriform glands). This promotes clinical adoption since AI-triage decisions can be linked to recognizable histomorphologic patterns. The impact of segmentation priors depends on mask quality and domain match. In prostate, where prior work has enabled the generation of accurate 3D gland segmentation masks ^18, 56^, segmentation guidance improves the accuracy of downstream tasks. For esophagus, a foundation segmentation model, BiomedParse ^32^, was used to generate gland segmentation masks. Although these masks were imperfect, SCOPE still achieved consistent gains, indicating that the framework can benefit from even approximate structural priors when combined with prototype pretraining and cross-slice context. This also suggests a clear pathway for further improvement as more robust, organ-specific and 3D-pathology-optimized segmentation models become available.

Despite the strong performance of SCOPE relative to prior methods, several limitations should be noted. First, while SCOPE is designed for label-scarce settings, the limited number of slice-level annotations may lead to suboptimal performance for rare morphologies and diagnostic categories. As the field of 3D pathology continues to mature and larger, more diverse datasets become available, we anticipate that the representational capacity and robustness of the SCOPE framework will further improve. Second, the segmentation priors used in this study are only for glandular vs. non-glandular tissue types. While this choice is well-matched to prostate grading and provides a natural structural unit for many glandular diseases, it may be less optimal for other diseases. For example, certain Gleason pattern 5 variants in prostate cancer, including single-cell or sheet-like growth patterns ^57, 58^, may not be optimally represented using gland-focused structural priors. Adapting SCOPE to other diseases will require selecting segmentation targets that are appropriate for the underlying disease biology and validating their utility for guiding prototyping. Finally, this study focused on the technical development of a highly performant pipeline for AI-triaged 3D pathology, but large-scale clinical studies are needed in the future to demonstrate the clinical benefits in terms of diagnostic accuracy and time savings for pathologists. It should also be noted that, since current treatment guidelines have been established based on conventional 2D histopathology, the future implementation of AI-triaged 3D pathology may require a re-calibration of treatment thresholds to account for the significantly increased detection sensitivity that would likely result from this new diagnostic paradigm.

In summary, SCOPE provides a framework for interpretable AI-assisted triage of volumetric pathology datasets. By generating depth-resolved risk profiles across intact specimens, SCOPE enables pathologists to prioritize diagnostically relevant regions within large 3D pathology volumes. More broadly, these results suggest that prototype-based AI triage may provide a clinically practical strategy for integrating 3D pathology into future diagnostic workflows.

## Methods

### Dataset description

We curated three large-scale 3D digital histopathology cohorts acquired using advanced open-top light-sheet microscopy (OTLS), including two prostate cohorts and one esophagus cohort. All 3D OTLS volumes were acquired in two channels (nuclei and cytoplasm) and subsequently false-colored to resemble standard hematoxylin and eosin (H&E) staining ^59^. The datasets were acquired using different generations of OTLS systems ^8–10^ with native submicron lateral resolution and axial resolution of approximately 3–4 *µ*m. Following acquisition, image tiles were stitched and fused using BigStitcher, and all volumes were downsampled by a factor of two. For downstream analysis, image patches were extracted and rescaled to an effective in-plane sampling of approximately 0.9 *µ*m/pixel, ensuring that patches from different cohorts represented comparable physical tissue regions.

#### Prostate cohorts

Two prostate datasets were used in this study: one from the University of Washington Medical Center (Prostate) and one from the University of Pennsylvania (Prostate2).

For the Prostate cohort, formalin-fixed, paraffin-embedded (FFPE) tissue blocks from 54 prostate cancer patients were collected. Each block corresponded to one of six anatomical regions routinely sampled during standard sextant (6-core) or 12-core prostate biopsy procedures. From each block, a simulated core-needle biopsy (approximately 1 mm *×* 1 mm *×* 15 mm) was extracted and imaged using a second-generation OTLS system ^8^. Among 112 imaged cores, 121 representative 2D cross-sections were selected and annotated by a panel of six pathologists. Consensus was determined by majority vote, with ties resolved by selecting the more severe (higher) Gleason Grade Group (GG). This yielded GG labels: GG1 (54 slices), GG2 (34), GG3 (5), and GG4–5 (28).

For the Prostate2 cohort, 59 FFPE prostatectomy specimens from 59 patients were processed into 3-mm-diameter punch biopsies (thickness *>* 0.5 mm) and imaged using a fourth-generation OTLS system ^10^. To establish the ground truth, three pathologists collaboratively reviewed the 2D cross-sections. For each sample, a single consensus GG label was assigned based on a joint evaluation of two neighboring slices spaced 60 *µ*m apart. The resulting labels included GG1 (23 slices), GG2 (31), GG3 (4), and GG4–5 (1).

#### Esophagus cohort

The Esophagus cohort included 125 biopsy and endoscopic mucosal resection (EMR) specimens from 29 patients collected at the University of Washington Medical Center and imaged with a third-generation OTLS system ^9^. A total of 503 2D cross-sections were annotated by three board-certified pathologists using a tiered consensus strategy. Each slice was initially reviewed by two pathologists; in cases of disagreement, a third pathologist acted as an adjudicator to determine the final diagnostic category. This process yielded the following labels: non-dysplastic (391 slices), low-grade dysplasia (71), high-grade dysplasia (26), and esophageal adenocarcinoma (15).

### Training pipeline

Let 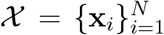 denote a cohort of *N* 3D OTLS volumes. Each volume **x**_*i*_ ∈ ℝ^*W ×H×Z*^ is a voxel grid of width *W*, height *H*, and depth *Z* (number of axial slices). We represent each volume as a stack of *x*–*y* planes, 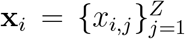, where *x*_*i,j*_ ∈ ℝ^*W ×H*^ denotes the *j*-th slice of the *i*-th specimen. For notational simplicity, channel dimensions are omitted.

Given the high-throughput nature of OTLS imaging, only a sparse subset of slices in each volume is annotated. Let *S*_*i*_ ⊆ *{*1, …, *Z}* denote the index set of labeled slices (SOIs) for volume *i*, and let *U*_*i*_ ⊆ *{*1, …, *Z}* denote the index set of unlabeled slices used for cohort-level pretraining (in this work, *U*_*i*_ includes all available unlabeled slices). Each labeled slice *x*_*i,j*_ (*j* ∈ *S*_*i*_) is assigned a diagnostic label *y*_*i,j*_ ∈ *{*1, …, *C}*, yielding the labeled dataset

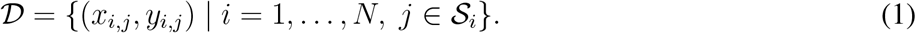

#### Multiple-instance learning

To accommodate computational constraints and the high dimensionality of x-y planes, we adopt a multiple-instance learning (MIL) framework commonly used in computational pathology. Under this paradigm, each slice *x*_*i,j*_ is treated as a *bag* consisting of many patches or *instances*. Specifically, each slice is partitioned into *N*_*i,j*_ non-overlapping patches 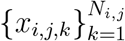, where 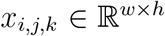 represents a patch of size *w × h*. A feature encoder *f*_enc_(*·*) maps each patch to a *d*_*e*_-dimensional embedding:

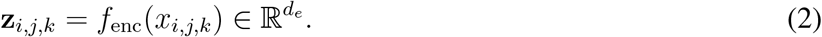

#### Attention-based aggregation

A widely adopted MIL aggregator is attention-based pooling ^36^. Given patch embeddings, attention weights *a*_*i,j,k*_ are computed as

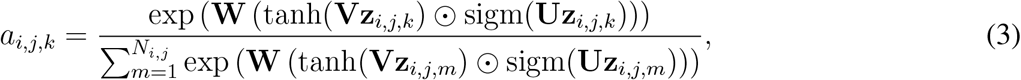

where **V**, 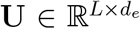 and **W** ∈ ℝ^1*×L*^ are learnable parameters, *L* is the attention hidden dimension, sigm(*·*) denotes the sigmoid function, and ⊙ denotes element-wise multiplication. The slice representation is

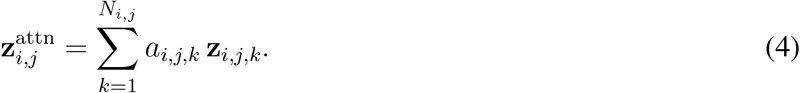

In label-scarce regimes, learning stable patch-level attention over thousands of instances can be challenging and may lead to overfitting, motivating a more compact aggregation strategy.

#### Prototype-based aggregation

SCOPE adopts a prototype-based MIL paradigm to obtain compact and interpretable slice representations. Given patch embeddings 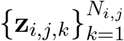, the key idea is to summarize a slice by a small vocabulary of recurring morphological prototypes rather than directly weighting a large number of individual patches. We model the distribution of patch embeddings using a Gaussian mixture model (GMM) with *M* components:

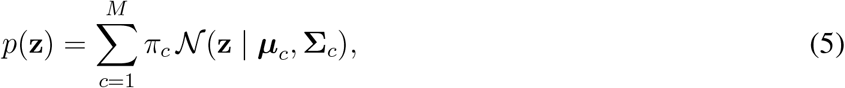

where each component *c* corresponds to one morphological prototype. Here, *π*_*c*_ ∈ ℝ is the *mixing weight* of prototype *c*, 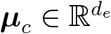 is its *mean* (prototype centroid), and 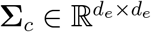 is its (diagonal) *covariance* describing within-prototype variability.

To fit the GMM parameters 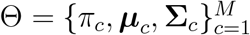, we use expectation–maximization (EM). In the E-step, we compute posterior responsibilities for all patch embeddings:

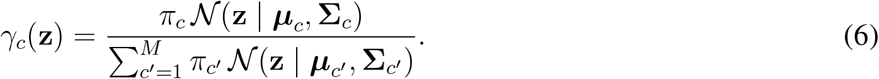

In the M-step, we update parameters:

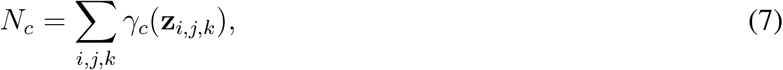

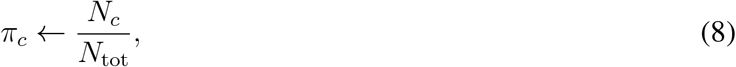

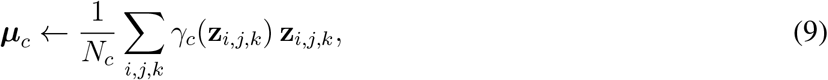

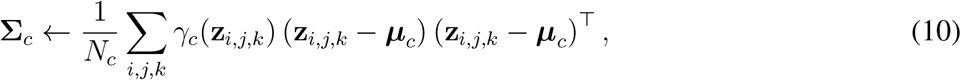

where 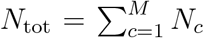 and 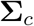 is constrained to be diagonal. In practice, we retain only the diagonal of the covariance update.

#### Prototype pretraining with unlabeled data

Directly fitting a GMM from random initialization can be unstable in high-dimensional embedding spaces. We therefore exploit the large amount of unlabeled 3D data to initialize the GMM means. Specifically, we pool patch embeddings from all unlabeled slices in the training cohort,

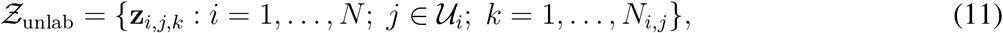

and run K-means clustering to obtain *M* centroids 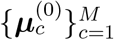. These centroids are used to initialize *{****µ*** *}* within the GMM model before EM fitting.

#### Segmentation-derived prior knowledge

To further enhance prototype quality, we incorporate domain-specific prior knowledge via tissue segmentation. This constrains prototyping to biologically meaningful tissue compartments and reduces the risk of mixing histologically distinct patterns that appear morphologically similar. For prostate cohorts, we utilize 3D-OTLS-gland-specific segmentation models ^18, 56^; for the esophagus cohort, we generate masks via the foundation model BiomedParse ^32^, as no task-specific model is currently available for this data.

Let *f*_seg_(*·*) denote a segmentation model that produces a volume-level discrete mask for each 3D pathology volume:

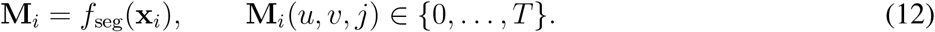

For slice *x*_*i,j*_, the corresponding mask is *M*_*i,j*_(*·, ·*) = **M**_*i*_(*·, ·, j*). For each patch, we crop a patch mask *m*_*i,j,k*_and compute the pixel-wise proportion 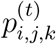 for each category *t* ∈ *{*1, …, *T}*. We assign a binary category tag

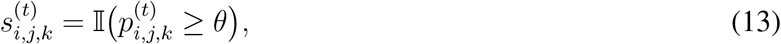

which defines segmentation-conditioned groups 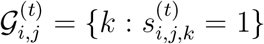. In this work, we set *T* = 2 (glandular and non-glandular) and learn separate prototype sets for the two compartments.

We then repeat the previously described K-means initialization *independently within each segmentation-defined tissue group*. For each group *t*, we run K-means on patch embeddings from unlabeled slices to obtain *M*_*t*_ initialization centroids 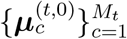.

#### Slice-specific GMM fitting

Given these group-specific centroids, we fit a slice-specific GMM within each tissue group using the EM algorithm. For each slice (*i, j*) and tissue group *t*, we estimate slice-specific

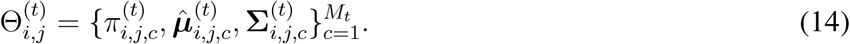

We initialize 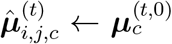 and 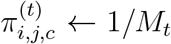, and set 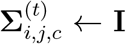 with diagonal covariance constraints in implementation. EM is run for a fixed number of 100 iterations using all patch embeddings from the slice.

Given the fitted parameters, the posterior responsibility of component *c* for a patch embedding **z** is

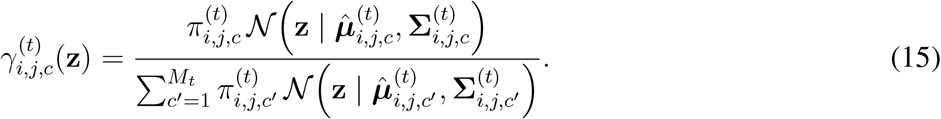

#### Slice-level prototype features

For each slice (*i, j*) and group *t*, we form a fixed-length group-specific feature by concatenating the slice-specific mixture weights and estimated component means:

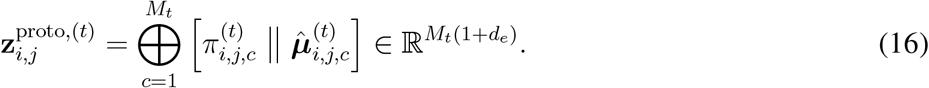

The final slice-level prototype feature is obtained by concatenating all group-specific features:

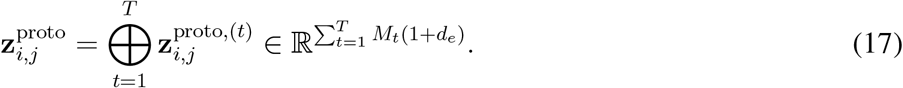

Although covariances 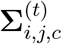 are estimated during EM and used when computing responsibilities, they are not included in 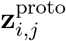 to reduce overfitting.

#### Cross-slice (2.5D) prototype aggregation

To leverage volumetric context, we incorporate depth-wise information from neighboring slices. For an SOI *x*_*i,j*_, we define a neighborhood of indices

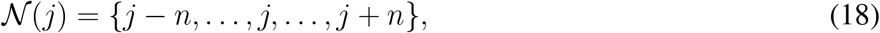

corresponding to a physical depth range Δ_*z*_ = 2*n d*_*z*_, where *d*_*z*_ is the axial spacing between sampled slices. For each neighboring slice *m* ∈ *N* (*j*) we compute 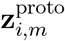 and fuse them using a lightweight attention module over slices:

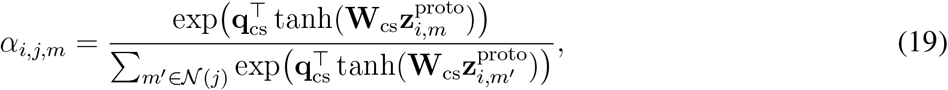

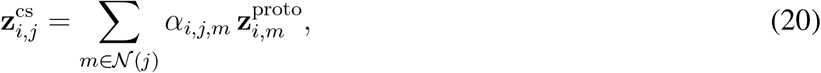

where **W**_cs_ and **q**_cs_ are learnable parameters for cross-slice aggregation.

By summarizing *N*_*i,j*_ patch embeddings into a compact prototype feature of dimensionality 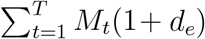, SCOPE substantially reduces the representation size compared with retaining all patch-level embeddings.

In general, SCOPE is trained in two phases.

##### Phase 1: cohort-level pretraining

We leverage all unlabeled slices across the training cohort to pretrain prototype centroids within each segmentation-defined tissue group. Concretely, for each group *t*, we pool patch embeddings 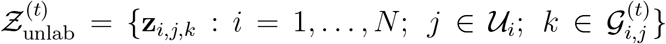 and run K-means to obtain initialization centroids 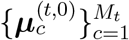.

##### Phase 2: slice-level training

Given the pretrained centroids, we fit the slice-specific GMM using the EM algorithm to compute 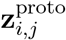 within each tissue group for each labeled SOI and its neighboring slices. We then fuse neighboring slices to obtain 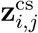 and train a linear classifier *f*_cls_(*·*) with cross-entropy loss:

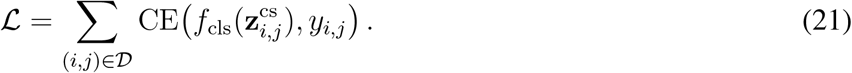

### Inference pipeline

At prediction time, the feature encoder *f*_enc_(*·*), the segmentation model *f*_seg_(*·*), the cross-slice attention module, and the classifier *f*_cls_(*·*) are fixed. For a given 3D volume **x**_*i*_, we first obtain a volume-level segmentation mask **M**_*i*_ and extract patch embeddings for each depth slice (and its neighboring slices) within the volume. For each slice index *j* and its neighborhood *N* (*j*), we fit the slice-specific GMM using the EM algorithm within each tissue group to compute slice-level prototype features 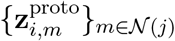. We then fuse neighboring-slice features with the trained cross-slice attention module to obtain 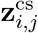 and apply *f*_cls_(*·*) to output a slice-level risk prediction. Repeating this over all slices yields a depth-resolved risk profile that is used to rank slices within each 3D volume for AI triage.

### Video Abstract

A video abstract summarizing the motivation, methodology, and key findings of this study accompanies this manuscript.

## Supporting information

Video Abstract

## Supplementary Videos

**Supplementary Video 1. Visualization of glandular prototype assignments in a representative prostate biopsy volume**.

A representative prostate biopsy volume from the Prostate cohort is shown. Each frame corresponds to an individual slice at a different imaging depth. Glandular regions are overlaid with their assigned morphological prototypes, illustrating how the learned prototypes capture distinct glandular architectures throughout the tissue volume.

**Supplementary Video 2. Depth-resolved risk prediction in a representative prostate biopsy volume**.

A representative prostate biopsy volume from the Prostate cohort is shown. Each frame corresponds to an individual slice at a different imaging depth. The risk score predicted by SCOPE is displayed in the left of each frame, illustrating how diagnostic risk varies across depth.

## Code availability

The source code used in this study is publicly available at https://github.com/RenaoYan/SCOPE.

## Author contributions

R. Yan, G. Gao, A.H. Song, F. Mahmood, and J.T.C. Liu conceived the study and designed the experiments. P. Lal, A. Madabbushi, L.D. True, and J.T.C. Liu collected clinical samples. G. Gao, D. Brenes, and S.S.L. Chow imaged and processed samples for the 3D pathology data cohorts. J. Shen, P. Lal, L.D. True, and D.M. Reddi reviewed the 3D pathology data. R. Yan and G. Gao ran all the computational experiments with guidance from A.H. Song. R. Yan, G. Gao, A.H. Song, H. Hsieh, Y. Zhao, C. Almagro-Pérez and J.T.C. Liu prepared the manuscript. All authors contributed to the writing. J.T.C. Liu supervised the research.

## Acknowledgements

This research was primarily supported by the National Institute of Diabetes and Digestive and Kidney Diseases (NIDDK) under R01DK138948 (Liu, Mahmood, and Grady) and the National Cancer Institute (NCI) under R01CA268207 (Liu and Madabhushi). NIH support for WM Grady includes U01CA152756, R01CA220004, U2CCA271902, U54CA163060, and U01CA182940. Funding for WM Grady is also provided by the Prevent Cancer Foundation, Cottrell Family Fund, Evergreen Fund, and Listwin Foundation to WMG. These studies are also supported by GiCaRes from UW Departments of Medicine (Gastroenterology Division) and Lab Medicine & Pathology. Additional NIH support includes grants R01EB031002 (Liu), R01CA268287 (Madabhushi), U01CA269181(Madabhushi), R01CA249992 (Madabhushi), R01CA202752 (Madabhushi), R01CA208236 (Madabhushi), R01CA216579 (Madabhushi), R01CA220581 (Madabhushi), R01CA257612 (Madabhushi), 1U01CA239055 (Madabhushi), 1U01CA248226 (Madabhushi), 1U54CA254566 (Madabhushi), R01HL15127701 (Madabhushi), R01HL15807101 (Madabhushi), and R01CA270437 (Shen). Support was also received from the VA Merit Review Award IBX004121A (Madabhushi) and sponsored research agreements from Bristol Myers-Squibb, and Astrazeneca. Additional support was through the Department of Defense (DoD) Prostate Cancer Research Program (PCRP) grant W81XWH-18-10358 (Liu and True), W81XWH-14-2-0183 (True), W81XWH-20-1-0851 (Madabhushi and Liu), the Pacific Northwest Prostate Cancer SPORE P50CA97186 (True), NSF Graduate Research Fellowship DGE-1762114 (K.W.B.), and the Canary Foundation. Research was also supported in part by the ARPA-H under contract number D24AC00357 (Liu) and D25AC00140 (Madabhushi). The content is solely the responsibility of the authors and does not necessarily represent the official views of the National Institutes of Health, the U.S. Department of Veterans Affairs, the Department of Defense, or the United States Government.

---

**Extended Data Figure 1:**
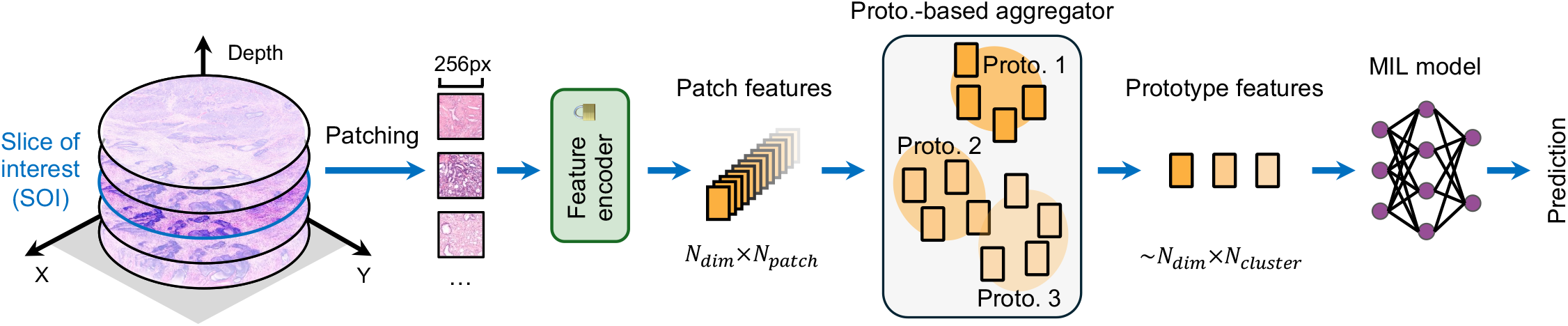
Prototype learning for AI triage. A slice of interest (SOI) is partitioned into patches and encoded into patch embeddings. Prototype-based aggregation then maps a large set of patch embeddings (size ~ *N*_patch_) to a compact prototype representation (size ~ *N*_proto_) by grouping morphologically similar patches into a small number of prototypes, yielding a fixed-length “prototype composition” feature for downstream tasks.

**Extended Data Figure 2:**
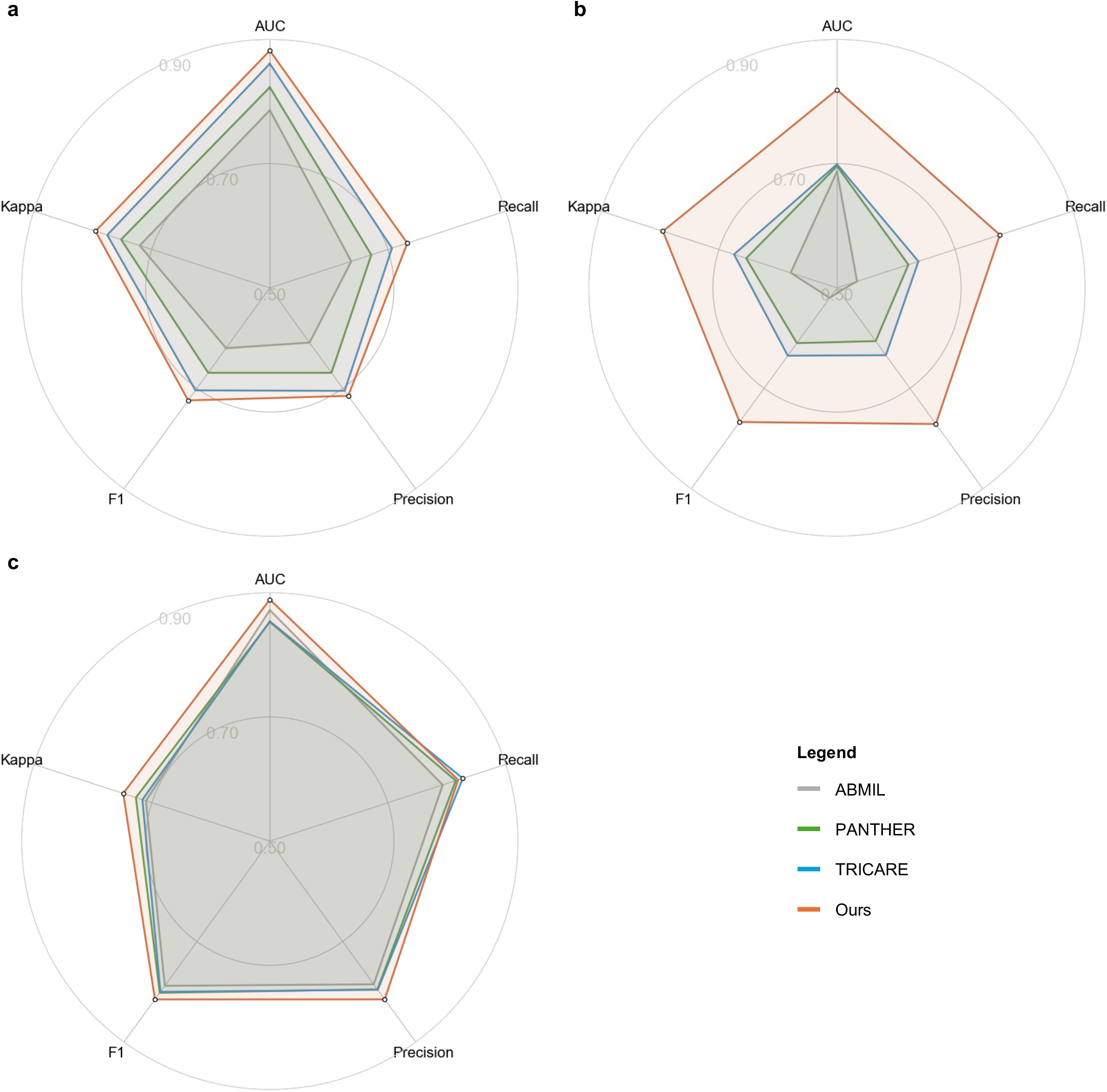
Radar-plot comparison of SCOPE and baselines across cohorts. Radar plots summarize performance across five metrics (AUC, precision, recall, F1, and Cohen’s *κ*), where larger radius indicates better performance. Values are averaged across leave-one-patient-out folds. Results are shown for **(a)** Prostate, **(b)** Prostate2, and **(c)** Esophagus cohorts.

**Extended Data Figure 3:**
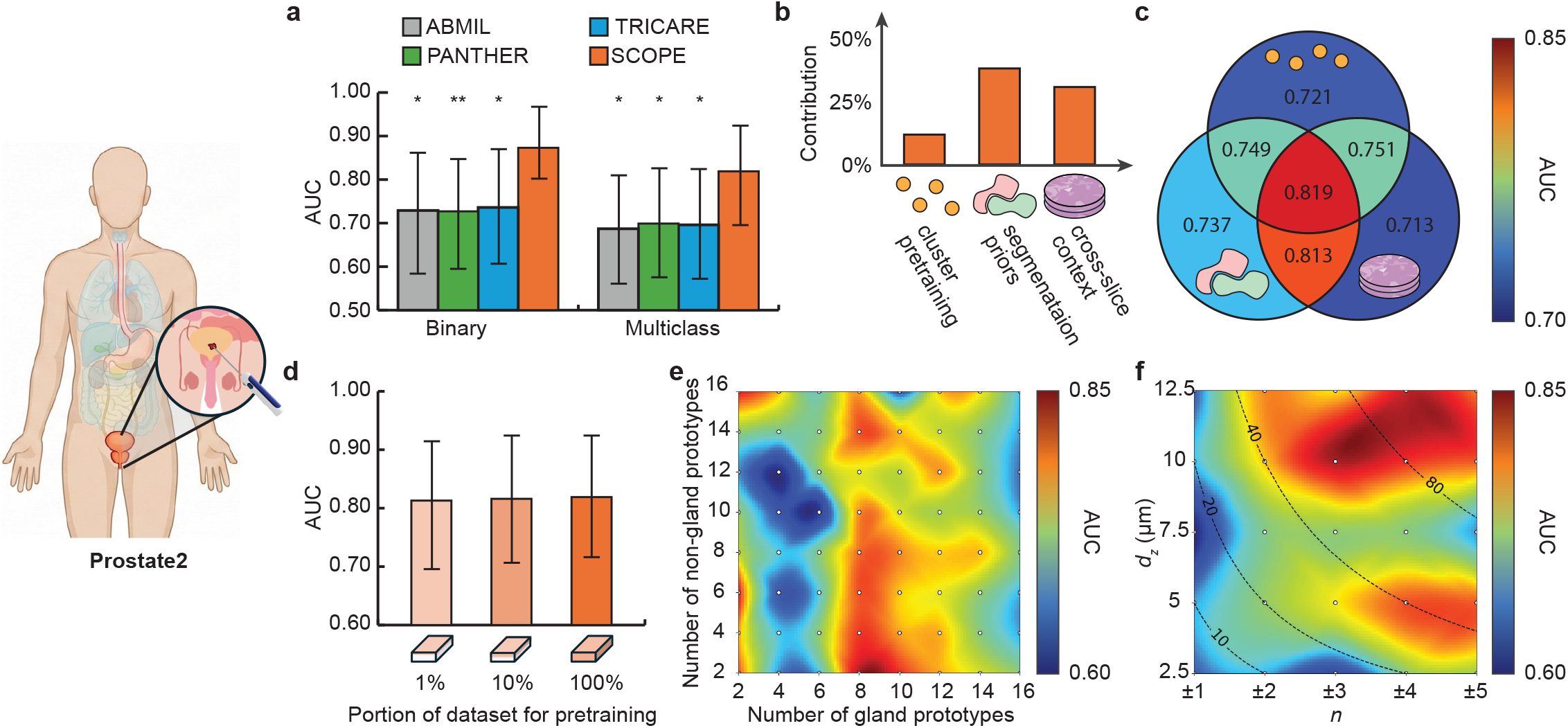
Risk assessment performance and ablations on the Prostate2 cohort. **(a)** Binary and multiclass AUC for SCOPE and baselines (ABMIL, PANTHER, TRICARE). **(b)** Shapley value (SHAP)-based contribution of the three design choices (prototype pretraining, segmentation priors, cross-slice context). **(c)** Performance of all 2^3^ component combinations (Venn diagrams; center = full SCOPE). **(d)** Effect of unlabeled pretraining scale on downstream AUC. **(e)** Sensitivity to the number of gland and non-gland prototypes. **(f)** Sensitivity to cross-slice context parameters (number of neighboring slices *n* and spacing *d*_*z*_, with Δ_*z*_ = *n × d*_*z*_). Bars show mean AUC across leave-one-patient-out folds; error bars indicate 95% confidence intervals. Statistical significance is shown for SCOPE versus each baseline (^*^*p* ≤ 0.05, ^**^*p* ≤ 0.01; nonparametric bootstrap test with 5,000 resamples).

**Extended Data Figure 4:**
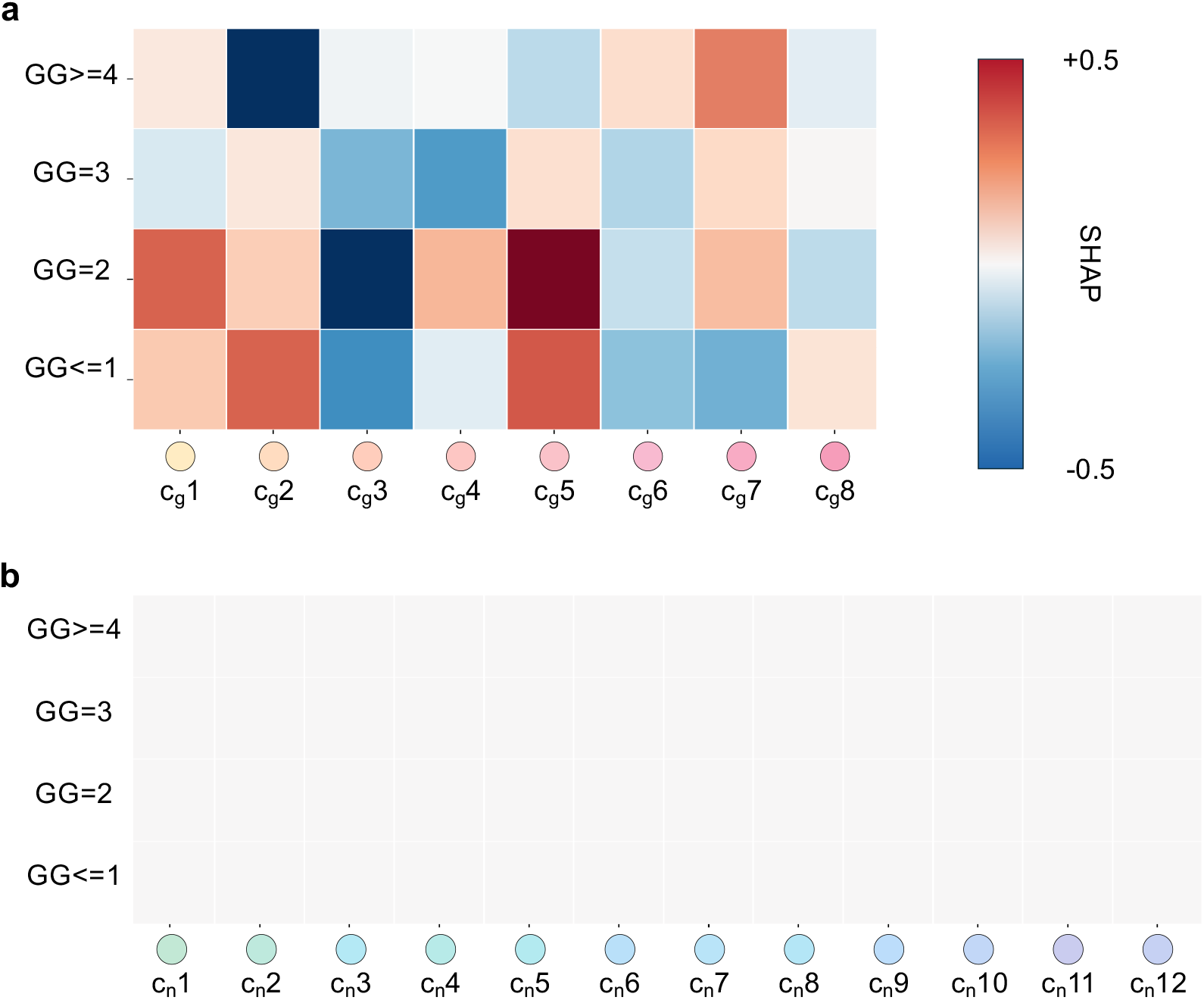
Prototype contribution analysis for Gleason grade group prediction (Prostate cohort). Heatmaps show Shapley values (SHAP) quantifying prototype-level contributions to multiclass grade-group predictions. Rows correspond to predicted grade groups (GG ≤ 1, GG2, GG3, GG ≥ 4), and columns correspond to learned prototypes within **(a)** glandular and **(b)** non-glandular compartments. Positive SHAP (red) indicates that the prototype increases the model score for that grade group, whereas negative SHAP (blue) indicates a decreasing effect. Glandular prototypes show both positive and negative associations with grade-group predictions, while non-glandular prototypes exhibit weak or near-zero contributions across classes, consistent with Gleason grading being driven primarily by glandular architecture.

**Extended Data Figure 5:**
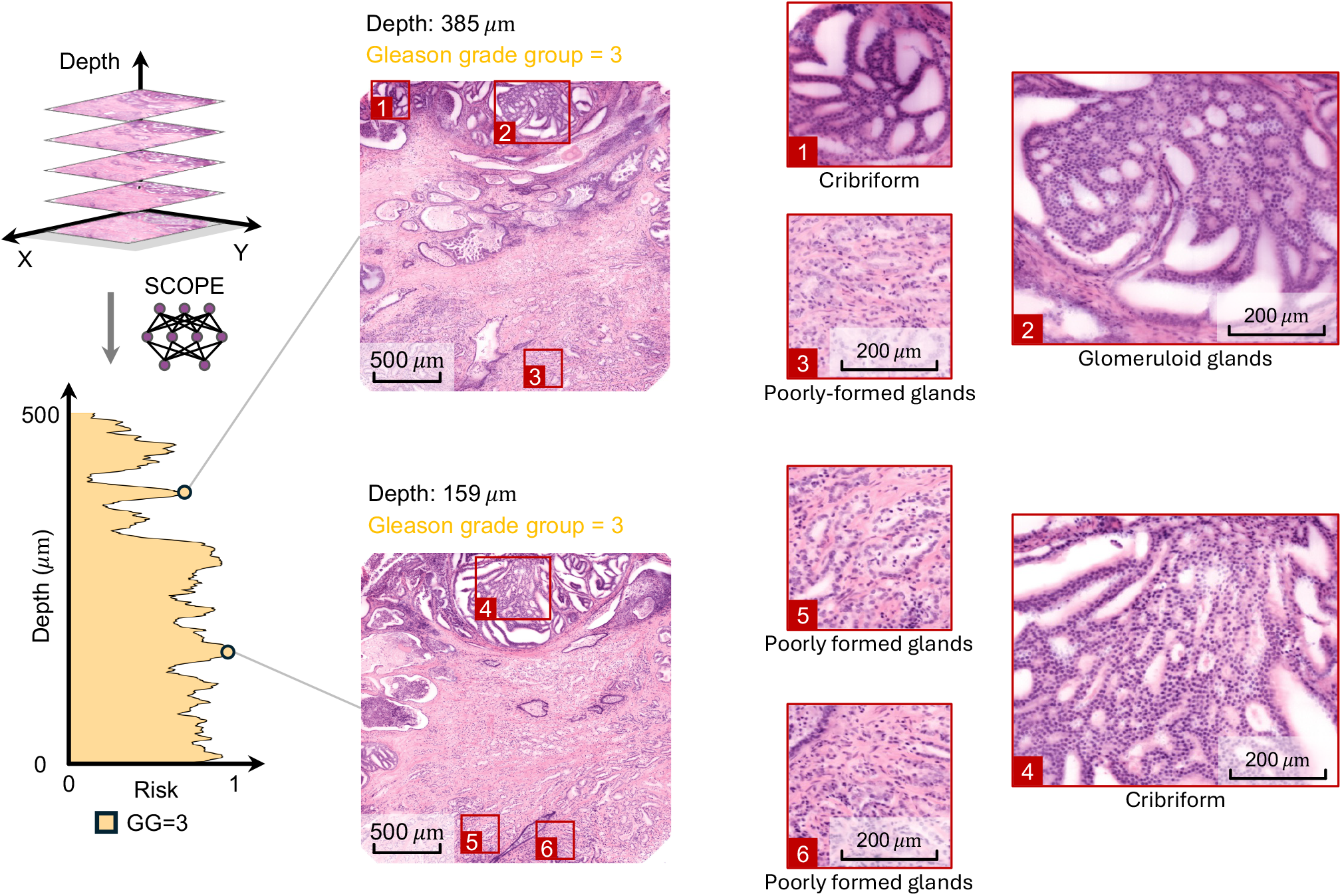
Interpretable 3D triage via risk–depth curves in representative Prostate2 biopsies. The left panel shows risk–depth curve generated by SCOPE across all depth slices; colors indicate the predicted class at each depth, and the curve height indicates the risk score (probability of the specimen-level target risk class). The right panel shows example whole-slice views at selected depths, with red boxes marking ROIs shown at higher magnification. Representative slices of different categories illustrate depth-dependent heterogeneity. Scale bars represent 500 *µ*m for whole-slice views, and 200 *µ*m for ROI zoom-ins.

## References

1. Siegel, R. L., Giaquinto, A. N. & Jemal, A. Cancer statistics, 2024. CA: A Cancer Journal for Clinicians 74, 12–49 (2024).

2. Stelzer, E. H. et al. Light sheet fluorescence microscopy. Nature Reviews Methods Primers 1, 73 (2021).

3. Ueda, H. R. et al. Whole-brain profiling of cells and circuits in mammals by tissue clearing and light-sheet microscopy. Neuron 106, 369–387 (2020).

4. Zhang, D. et al. Spatial analysis of tissue immunity and vascularity by light sheet fluorescence microscopy. Nature protocols 19, 1053–1082 (2024).

5. Patel, K. B. et al. High-speed light-sheet microscopy for the in-situ acquisition of volumetric histological images of living tissue. Nature biomedical engineering 6, 569–583 (2022).

6. Park, W. Y. et al. Open-top bessel beam two-photon light sheet microscopy for three-dimensional pathology. Elife 12, RP92614 (2024).

7. Chen, L. et al. Efficient 3d imaging and pathological analysis of the human lymphoma tumor microenvironment using light-sheet immunofluorescence microscopy. Theranostics 14, 406 (2024).

8. Glaser, A. K. et al. Multi-immersion open-top light-sheet microscope for high-throughput imaging of cleared tissues. Nature communications 10, 2781 (2019).

9. Barner, L. A., Glaser, A. K., Huang, H., True, L. D. & Liu, J. T. Multi-resolution open-top light-sheet microscopy to enable efficient 3d pathology workflows. Biomedical Optics Express 11, 6605–6619 (2020).

10. Glaser, A. K. et al. A hybrid open-top light-sheet microscope for versatile multi-scale imaging of cleared tissues. Nature methods 19, 613–619 (2022).

11. Bishop, K. W. et al. Axially swept open-top light-sheet microscopy for densely labeled clinical specimens. Optics Letters 49, 3794–3797 (2024).

12. Bishop, K. W. et al. An end-to-end workflow for nondestructive 3d pathology. Nature Protocols 19, 1122–1148 (2024).

13. Baraznenok, E. et al. Assessing the effects of a 3d pathology tissue-processing workflow on downstream molecular analyses. bioRxiv 2026–02 (2026).

14. Farahani, N. et al. Three-dimensional imaging and scanning: current and future applications for pathology. Journal of pathology informatics 8, 36 (2017).

15. Liu, J. T. et al. Harnessing non-destructive 3d pathology. Nature biomedical engineering 5, 203–218 (2021).

16. Liu, J. T. et al. Engineering the future of 3d pathology. The Journal of Pathology: Clinical Research 10, e347 (2024).

17. Liu, J. T. et al. Engineering the future of 3d pathology. The Journal of Pathology: Clinical Research 10, e347 (2024).

18. Xie, W. et al. Prostate cancer risk stratification via nondestructive 3d pathology with deep learning–assisted gland analysis. Cancer research 82, 334–345 (2022).

19. Serafin, R. et al. Nondestructive 3d pathology with analysis of nuclear features for prostate cancer risk assessment. The Journal of Pathology 260, 390–401 (2023).

20. Song, A. H. et al. Analysis of 3d pathology samples using weakly supervised ai. Cell 187, 2502–2520 (2024).

21. Erion Barner, L. A. et al. Artificial intelligence-triaged 3-dimensional pathology to improve detection of esophageal neoplasia while reducing pathologist workloads. Modern pathology: an official journal of the United States and Canadian Academy of Pathology, Inc 36, 100322 (2023).

22. Gao, G. et al. Triage of 3d pathology data via 2.5 d multiple-instance learning to guide pathologist assessments. In Proceedings of the IEEE/CVF Conference on Computer Vision and Pattern Recognition, 6955–6965 (2024).

23. Gao, G. et al. Deep-learning triage of 3d pathology datasets for comprehensive and efficient pathologist assessments. bioRxiv (2025).

24. Song, A. H. et al. Artificial intelligence for digital and computational pathology. Nature Reviews Bioengineering 1, 930–949 (2023).

25. Song, A. H. et al. Morphological prototyping for unsupervised slide representation learning in computational pathology. In Proceedings of the IEEE/CVF Conference on Computer Vision and Pattern Recognition, 11566–11578 (2024).

26. Yao, J., Zhu, X., Jonnagaddala, J., Hawkins, N. & Huang, J. Whole slide images based cancer survival prediction using attention guided deep multiple instance learning networks. Medical image analysis 65, 101789 (2020).

27. Vu, Q. D., Rajpoot, K., Raza, S. E. A. & Rajpoot, N. Handcrafted histological transformer (h2t): Unsupervised representation of whole slide images. Medical image analysis 85, 102743 (2023).

28. Claudio Quiros, A. et al. Mapping the landscape of histomorphological cancer phenotypes using self-supervised learning on unannotated pathology slides. Nature Communications 15, 4596 (2024).

29. Cutler, K. J. et al. Omnipose: a high-precision morphology-independent solution for bacterial cell segmentation. Nature methods 19, 1438–1448 (2022).

30. Pang, M., Roy, T. K., Wu, X. & Tan, K. Cellotype: a unified model for segmentation and classification of tissue images. Nature methods 22, 348–357 (2025).

31. Isensee, F., Jaeger, P. F., Kohl, S. A., Petersen, J. & Maier-Hein, K. H. nnu-net: a self-configuring method for deep learning-based biomedical image segmentation. Nature methods 18, 203–211 (2021).

32. Zhao, T. et al. A foundation model for joint segmentation, detection and recognition of biomedical objects across nine modalities. Nature methods 22, 166–176 (2025).

33. Ma, J. et al. Segment anything in medical images. Nature Communications 15, 654 (2024).

34. Stringer, C., Wang, T., Michaelos, M. & Pachitariu, M. Cellpose: a generalist algorithm for cellular segmentation. Nature methods 18, 100–106 (2021).

35. Almagro-Pérez, C. et al. Ai-driven 3d spatial transcriptomics. arXiv preprint arXiv:2502.17761 (2025).

36. Ilse, M., Tomczak, J. & Welling, M. Attention-based deep multiple instance learning. In International conference on machine learning, 2127–2136 (PMLR, 2018).

37. Yufei, C. et al. Bayes-mil: A new probabilistic perspective on attention-based multiple instance learning for whole slide images. In The Eleventh International Conference on Learning Representations (2022).

38. Campanella, G. et al. Clinical-grade computational pathology using weakly supervised deep learning on whole slide images. Nature medicine 25, 1301–1309 (2019).

39. Chen, R. J. et al. Scaling vision transformers to gigapixel images via hierarchical self-supervised learning. In Proceedings of the IEEE/CVF Conference on Computer Vision and Pattern Recognition, 16144–16155 (2022).

40. Li, B., Li, Y. & Eliceiri, K. W. Dual-stream multiple instance learning network for whole slide image classification with self-supervised contrastive learning. In Proceedings of the IEEE/CVF conference on computer vision and pattern recognition, 14318–14328 (2021).

41. Lu, M. Y. et al. Data-efficient and weakly supervised computational pathology on whole-slide images. Nature biomedical engineering 5, 555–570 (2021).

42. Shao, Z. et al. Transmil: Transformer based correlated multiple instance learning for whole slide image classification. Advances in neural information processing systems 34, 2136–2147 (2021).

43. Zhu, L. et al. An accurate prediction of the origin for bone metastatic cancer using deep learning on digital pathological images. EBioMedicine 87 (2023).

44. Zhang, H. et al. Dtfd-mil: Double-tier feature distillation multiple instance learning for histopathology whole slide image classification. In Proceedings of the IEEE/CVF Conference on Computer Vision and Pattern Recognition, 18802–18812 (2022).

45. Javed, S. A. et al. Additive mil: intrinsically interpretable multiple instance learning for pathology. Advances in Neural Information Processing Systems 35, 20689–20702 (2022).

46. Zhu, L. et al. Hierarchically optimized multiple instance learning with multi-magnification pathological images for cerebral tumor diagnosis. IEEE Journal of Biomedical and Health Informatics (2025).

47. Yan, R. et al. Shapley values-enabled progressive pseudo bag augmentation for whole-slide image classification. IEEE transactions on medical imaging 44, 588–597 (2024).

48. Zhang, Y. et al. Attention-challenging multiple instance learning for whole slide image classification. arXiv preprint arXiv:2311.07125 (2023).

49. Shao, D. et al. Do multiple instance learning models transfer? In International Conference on Machine Learning, 54219–54238 (PMLR, 2025).

50. Tang, W. et al. Feature re-embedding: Towards foundation model-level performance in computational pathology. In Proceedings of the IEEE/CVF conference on computer vision and pattern recognition, 11343–11352 (2024).

51. Li, J. et al. Dynamic graph representation with knowledge-aware attention for histopathology whole slide image analysis. In Proceedings of the IEEE/CVF conference on computer vision and pattern recognition, 11323–11332 (2024).

52. Lu, M. Y. et al. A visual-language foundation model for computational pathology. Nature Medicine 30, 863–874 (2024).

53. Chen, R. J. et al. Towards a general-purpose foundation model for computational pathology. Nature medicine 30, 850–862 (2024).

54. Dempster, A. P., Laird, N. M. & Rubin, D. B. Maximum likelihood from incomplete data via the em algorithm. Journal of the royal statistical society: series B (methodological) 39, 1–22 (1977).

55. Shapley, L. S. et al. A value for n-person games (1953).

56. Wang, R. et al. Direct three-dimensional segmentation of prostate glands with nnu-net. Journal of biomedical optics 29, 036001–036001 (2024).

57. Epstein, J. I. et al. The 2014 international society of urological pathology (isup) consensus conference on gleason grading of prostatic carcinoma: definition of grading patterns and proposal for a new grading system. The American journal of surgical pathology 40, 244–252 (2016).

58. Epstein, J. I. et al. A contemporary prostate cancer grading system: a validated alternative to the gleason score. European urology 69, 428–435 (2016).

59. Serafin, R., Xie, W., Glaser, A. K. & Liu, J. T. Falsecolor-python: a rapid intensity-leveling and digital-staining package for fluorescence-based slide-free digital pathology. Plos one 15, e0233198 (2020).

